# Loss of function in *RBBP5* results in a syndromic neurodevelopmental disorder associated with microcephaly

**DOI:** 10.1101/2024.02.06.578086

**Authors:** Yue Huang, Kristy L. Jay, Alden Yen-Wen Huang, Jijun Wan, Sharayu V. Jangam, Odelia Chorin, Annick Rothschild, Ortal Barel, Milena Mariani, Maria Iascone, Han Xue, Undiagnosed Diseases Network, Jing Huang, Cyril Mignot, Keren Boris, Virginie Saillour, Annelise Y. Mah-Som, Stephanie Sacharow, Farrah Rajabi, Carrie Costin, Shinya Yamamoto, Oguz Kanca, Hugo J. Bellen, Jill A. Rosenfeld, Christina G S Palmer, Stanley F. Nelson, Michael F. Wangler, Julian A Martinez-Agosto

## Abstract

**Purpose:** Epigenetic dysregulation has been associated with many inherited disorders. *RBBP5* encodes a core member of the protein complex that methylates histone 3 lysine-4 (H3K4) and has not been implicated in human disease.

**Methods:** We identify five unrelated individuals with *de novo* heterozygous pathogenic variants in *RBBP5*. Three truncating and two missense variants were identified in probands with neurodevelopmental symptoms including global developmental delay, intellectual disability, microcephaly, and short stature. Here, we investigate the pathogenicity of the variants through protein structural analysis and transgenic *Drosophila* models.

**Results:** Both missense p.T232I and p.E296D variants affect evolutionarily conserved amino acids and are expected to interfere with the interface between RBBP5 and the histones. In *Drosophila,* ubiquitous overexpression of human *RBBP5* is lethal in the larval developmental stage. Loss of *Rbbp5* leads to a reduction in brain size, and the human reference, p.T232I, or p.E296D variant transgenes fail to rescue loss of *Rbbp5.* Expression of either missense variant in an *Rbbp5* null background results in a less severe microcephaly phenotype than the human reference, indicating both p.T232I and p.E296D variants are loss-of-function alleles.

**Conclusion:** *De novo* heterozygous variants in *RBBP5* are associated with a syndromic neurodevelopmental disorder.

**Graphical abstract:** 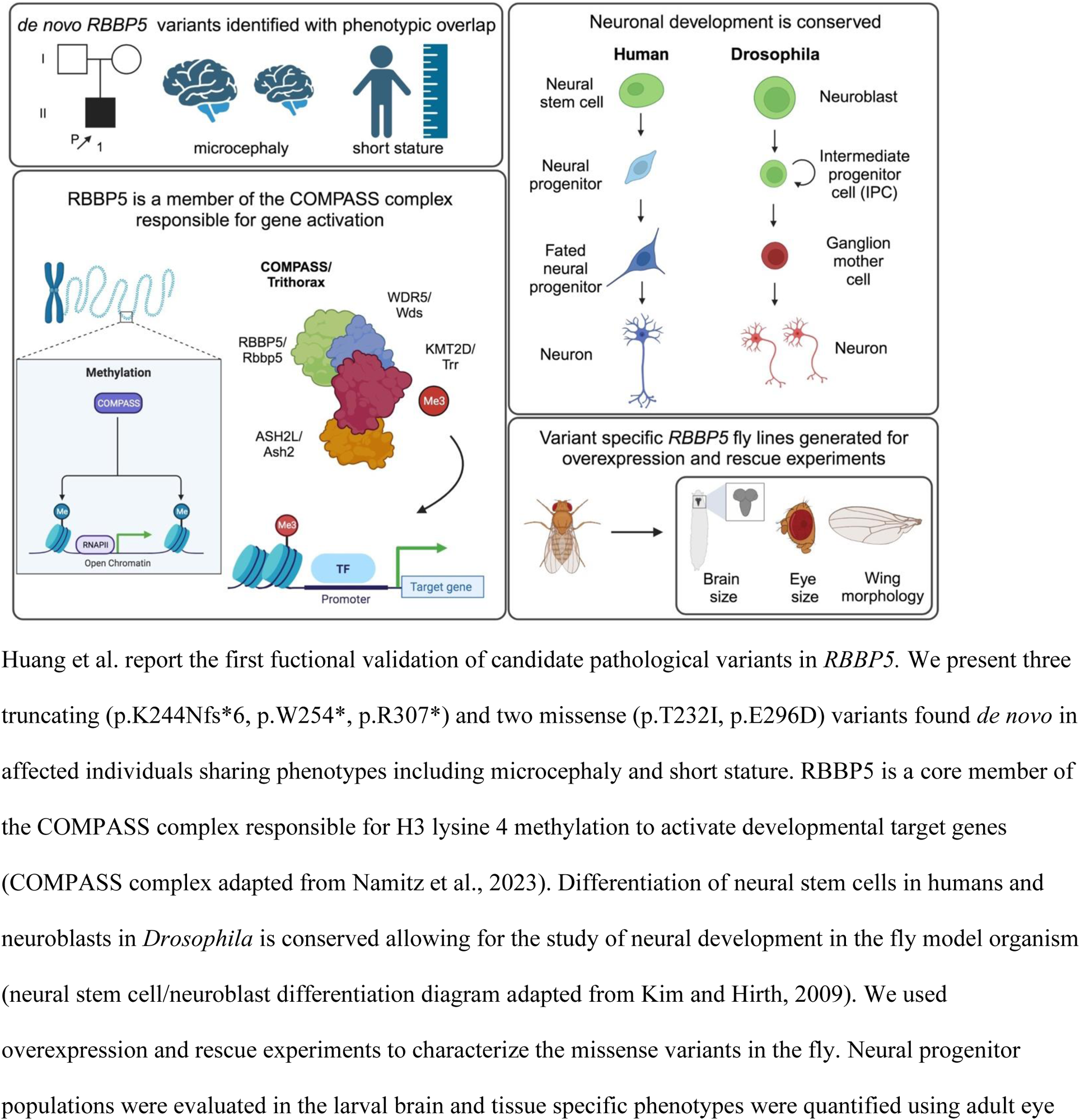
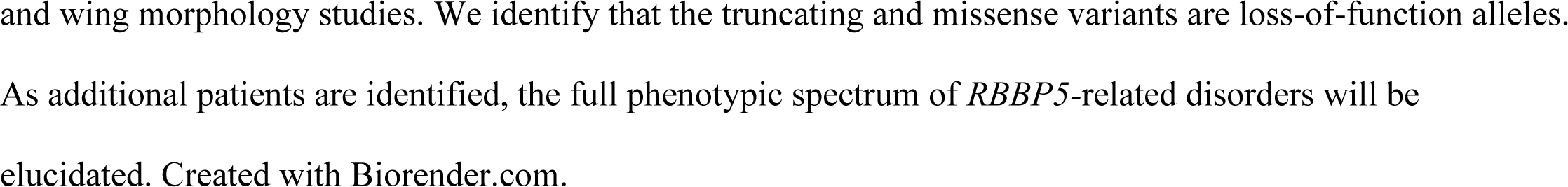

## INTRODUCTION

The epigenetic machinery has an essential role in the spatiotemporal regulation of gene expression. One of the main epigenetic regulatory mechanisms is the posttranslational modifications of histones, including methylation, acetylation, phosphorylation, and ubiquitylation^1^. These histone modifications allow precise and dynamic regulation of the accessibility of genomic regions to DNA-dependent processes including transcription, replication, DNA repair, and recombination^2^. The methylation of histone 3 lysine 4 (H3K4) is an evolutionarily conserved chromatin mark that is typically found in active transcription sites and is considered a marker for gene activation^3^. H3K4 methylation is predominantly mediated by the COMPASS (COMplex of Proteins Associated with Set1) protein complex, which includes one of the six SET1 domain-containing methyltransferases (KMT2A-F) and three other core members WDR5, ASH2L and RBBP5 to modulate the catalytic activity of methyltransferases^4^.

The number of Mendelian disorders caused by disruption of epigenetic machinery have greatly expanded in the past decade^5^. The functional classification divides the genes into four groups: writer, eraser, reader and remodeler^5^. Various histone methylation writers, including the SET1 domain-containing methyltransferases in COMPASS, have been associated with genetic disorders such as Kabuki syndrome (MIM#147920)^6^. However, the non-methyltrasnferase core members of COMPASS have not been linked to a human disorder to date. A heterozygous pathogenic variant in *KMT2D* accounts for 50-70% cases with Kabuki syndrome, and 20-30% of patients with a clinical diagnosis of Kabuki syndrome have no identified known variant that causes the disease^6^. It has been hypothesized that a pathogenic mutation in other core members of the COMPASS protein complex could contribute to cases with Kabuki-like phenotypes^7^.

In this study, we report five unrelated patients with *de novo* heterozygous variants in *RBBP5*. These individuals present with neurodevelopmental features including intellectual disability, developmental delay, microcephaly, and short stature. We provide evidence to support the pathogenicity of the variants through bioinformatic analysis, protein structure modeling, and functional studies in *Drosophila.* The difficulty in diagnosing Individual 1 led to his enrollment in the Undiagnosed Diseases Network (UDN). The goal of the UDN is to facilitate collaboration between clinicians and researchers to improve diagnosis and care^8–10^. In addition to taking advantage of state-of-the-art phenotyping and genotyping tools, the UDN uses model organisms such as *Drosophila melanogaster* to perform functional assays on rare genetic variants identified in rare disease patients^11,12^. Fly researchers have generated an extensive transgenic toolkit for *Drosophila* allowing for rapid functional characterization of candidate pathological variants^13–16^. Functional validation of genetic variants is a critical step toward confirming diagnosis, and efforts to characterize novel disease genes improve diagnostic success of additional patients in the future^12,17–20^. Elucidating the mechanism for rare and undiagnosed diseases leads to improved diagnostic rates, earlier intervention, possible targeted therapeutics, and ultimately improved quality of life^20^.

## MATERIALS AND METHODS

### Identification of individuals

Individual 1 was referred to the Undiagnosed Disease Network at UCLA by a local physician. Other individuals were identified in GeneMatcher^21^, and clinical information were collected through collaborators. In addition, a thorough search for candidate *RBBP5* variants was conducted in the clinical exome/genome database at the Baylor Genetics Laboratories. All individuals provided written consent for participation of research and publication, including consent to publish patient photos. This study has been approved by the institution review board at UCLA. *RBBP5* variants were identified by either exome or genome sequencing. All individuals except one had both biological parents as comparators in the sequencing and the variants were confirmed to be *de novo* due to absence in the parental sequences. Sanger sequencing was not performed because of high quality of variant calling.

### Structural analysis

Structural analysis of human MLL3-ubNCP complex (PDB: 6KIW) was carried out with PyMOL (The PyMOL Molecular Graphics System, Version 2.0 Schrödinger, LLC, https://pymol.org/2/)

#### Drosophila melanogaster

Fly lines were obtained from the Bloomington Drosophila stock center and the Kyoto Drosophila stock center: daughterless-GAL4(3) (*daughterless^GAL4^*) (Bellen lab), w[*]; P{w[mC]=UAS-mCherry.NLS}3 (BDSC: 38424) (*UAS-mCherry.NLS*), y[1]w[*]; P{w[+mC]=Act5C-GAL4}25Fo1/Cyo, y[+] (BDSC:4414) (*Actin^GAL4^*), y[1]w[1118]; P{w[+mC]=ey3.5-GAL4.Exel}2 (BDSC: 8220) (*eyeless^GAL4^*), P{w[+mC]=GAL4-elav.L}2/Cyo (BDSC: 8765) (*elav^GAL4^*), w[1118]; P{w[+m*]=GAL4}repo/TM3, Sb (BDSC: 7415) (*repo^GAL4^*), UAS-lacZ (Bellen lab) w[*];Cyo, P{w[+mC]=Tb[1]Cpr[Cyo-A]/sna[Sco] (BDSC: 36335) (*Cyo, Tb*), w[1118]; Df(3L)BSC447/TM6C, Sb[1] cu[1] (BDSC:24951) (*Df (3L)BSC447*), y^1^ w*; PBac{y^+mDint2^ w^+mC^=UAS-hsp-Hsap\RBBP5.HA.1}VK00037 / CyO, P{ry^+t7.2^=sevRas1.V12}FK1 (DGRC: 305207) (*RBBP5-HA*). Stocks were held at 25°C and all experiments were carried out at 25°C unless specified.

### *Rbbp5^Kozak GAL4^* transgenic line generation

The *Rbbp5^Kozak^ ^GAL4^* was generated as previously described (Kanca et al., 2022). Briefly, the *Rbbp5* coding sequence was replaced with a *Kozak sequence-GAL4-polyA-FRT-3XP3EGFP-polyA-FRT* (*KozakGAL4*) cassette using CRISPR mediated homologous recombination. The homology donor intermediate construct is prepared by synthesis of sgRNA targeting the 5’ and 3’ untranslated regions (AAAATGAATTTGGAGCTACTAGG and TTATTTCTTGGTACGTCCGGCGG respectively) and short (200 bp) homology arms in pUC57_Kan_gw_OK2 vector. KozakGAL4-polyA-FRT-3XP3EGFP-FRT cassette is subcloned in the homology donor intermediate to generate the homology donor construct. Homology donor construct is injected (250 ng/µl) in embryos containing Cas9 in their germline (*y^1^w*; attP40(y+){nos-Cas9(v+)}; iso5)* and the resulting G0 progeny are crossed to *y^1^w\** flies and screened for *3XP3-EGFP* expression. The correct integration of KozakGAL4 cassette in the proper locus is verified by PCR using forward and reverse primers flanking the homology arms and construct specific forwards and reverse primers as described in Kanca et al. 2022. The *RBBP5^Kozak^ ^GAL4^* transgenic line was submitted to the Bloomington Drosophila Stock Center (BDSC: 97331).

### Human *RBBP5* and *Drosophila Rbbp5* construct generation

Human pcDNA3-FLAG-RBBP5 plasmid used in the protein expression experiments was obtained from Addgene (Cat# 15550, MA, USA). Candidate variants were introduced using QuikChange II Site-directed mutagenesis kit according to manufacturer’s protocol (Agilent, CA, USA). The mutations were confirmed by Sanger sequencing. For *Drosophila* experiments, Human cDNA for *RBBP5* (clone: IOH28957) was obtained from the collection of the late Dr. Kenneth Scott at Baylor College of Medicine. Q5 site directed mutagenesis was completed to create the variant sequence from the reference (c.695C>T, p.T232I, c.888A>T, p.E296D). Constructs were transformed using high efficiency *E. coli* competent cells (New England Biolabs, MA, USA, Cat # C2987H) from the pDONR 221 entry vector to the pGW-HA.attB destination vector using Gateway cloning and the sequences were confirmed by Sanger sequencing. Vectors were injected into embryos to create *P{UASt-RBBP5-Ref}VK37, P{UASt-RBBP5-T232I}VK37,* and *P{UASt-RBBP5-E296D}VK37*. For fly cDNA construct generation, *Rbbp5* (NM_140952.3) wild-type and variant (p.T231I and p.E295D) lines were obtained (clone OFa19095D, GenScript USA, Inc., NJ, USA). Constructs were transformed using high efficiency *E. coli* competent cells from the pGenDONR vector to the Gateway compatible pDONR 223 entry vector and subcloned to the pGW-HA.attB destination vector. Sequences were confirmed using Sanger sequencing and vectors were injected into embryos to create *P{UASt-Rbbp5}VK37, P{UASt-Rbbp5-T231I}VK37,* and *P{UASt-RBBP5-E295D}VK37.* Transgenic males are crossed to *y^1^ w\** stocks and construct integration is confirmed through selection for the mini white gene.

### RBBP5 protein expression and Western blot

Flag-tagged wild type and mutated human *RBBP5* plasmid were transfected to HEK293 cells by lipofectamine 3000 according to manufacturer’s protocol (Invitrogen, CA, USA). Cells were lysed by RIPA buffer and 20μg of whole cell lysate were used to assess protein expression in Western blot. ANTI-FLAG® M2 antibody from Sigma and anti-beta-actin from Santa Cruz Biotechnology were used to detect FLAG-tagged RBBP5 and Beta-actin respectively. For *Drosophila* experiments, histone extraction was performed (Abcam, Cambridge, UK, Cat# Ab113476) with five whole 3^rd^ instar larvae (n=8 replicates) using the manufacturer’s protocol. Protein (10 μL) was loaded of each sample and RBBP5 (Cell Signaling Technology (CST), MA, USA Cat#12766), H3K4me3 (CST, MA, USA Cat# 9733), and histone H3 (CST, MA, USA Cat# 9715) primary antibodies were used with Goat anti-Rabbit HRP and imaged on a Bio-Rad Chemidoc MP imaging system. H3K4me3 normalization to total H3 was completed using ImageJ.

### *RBBP5/Rbbp5* overexpression viability and morphology evaluation in *Drosophila*

Heterozygous or homozygous (in the case of larval and pupal lethal crosses) male UAS-cDNA stocks were crossed to female GAL4 ubiquitous (*Actin^GAL4^, daughterless^GAL4^*) and tissue specific (*eyeless^GAL4^, elav^GAL4^, repo^GAL4^*) lines. The resulting progeny were counted by genotype (n>50). Viability was calculated by comparing the observed number progeny to the expected (o/e ratio) based on Mendelian inheritance patterns. A normalized o/e ratio greater than 0.8 is defined as viable, semi-lethality is 0.8-0.15, and lethality is less than 0.15 and the latest developmental stage observed is reported as the lethal point. For *da^GAL4^* overexpression, only latest developmental stage reached is reported since the *da^GAL4^* stock is not balanced. Overexpression based phenotypes are scored in greater than five larvae or adult flies.

### *RBBP5* human cDNA rescue evaluation

In human cDNA rescue experiments, *Rbbp5^Kozak^ ^GAL4^* lethality is reported as the latest developmental stage reached. Non-balanced dead pupae (n=19/341) were observed in self-crosses of *Rbbp5^Kozak^ ^GAL4^ / TM6B,Sb,Tb* stocks and dead larvae were also observed in the crosses. Therefore, there is the possibility that *Rbbp5^Kozak^ ^GAL4^* is lethal at the embryo, larval, and pupal stages, however for the purposes of this study, rescue was evaluated as the ability of the human cDNA to rescue to the adult developmental stage in (n>40) progeny.

### *Drosophila* developmental staging

The following characteristics were selected to stage animals: early L3 larvae have branched but not extruding spiracles and developed posterior spiracles; and late L3 larvae have extruding spiracles, visible gut clearance, and exhibit wandering behavior.

### Brain immunostaining and brain lobe quantification

For larval counterselection, homozygous *da^GAL4^* lines were used and *Actin^GAL4^* was crossed to *Cyo, Tb* to create *Actin-GAL4 / Cyo,Tb.* These lines were then crossed to homozygous *RBBP5^Ref^, RBBP5^T232I^* and *RBBP5^E296D^* stocks. 3^rd^ instar larvae (gut clearance, branched spiracles, wandering behavior) were dissected in ice-cold PBS. Larval brain preps were fixed in 4% PFA/PBS/2% Triton overnight. Brains were blocked in PBS/2%Triton/5% normal donkey serum for 1 hour. Primary antibodies were incubated overnight; rat anti-Deadpan (Abcam, Cambridge, UK, Cat# ab195173, 1:250) and mouse anti-Prospero (Developmental Studies Hybridoma Bank, IA, USA Cat# MR1A, 1:1000). Secondary antibodies were incubated for 2 hours at room temperature; rat anti-GFP and mouse anti-Cy3 (1:250). Brains were mounted and imaged using a Zeiss 710 confocal microscope (Neurovisualization core, Baylor College of Medicine, TX, USA) with 1 μm sections. The area of one brain lobe was quantified using the area tool in Image J.

### Eye area quantification

Whole heads were imaged using a Leica KL1500 LCD microscope using 10x magnification. The area of one eye (n=5) was quantified using the area tool in ImageJ.

### Statistical analysis

Statistical anlayis was completed using GraphPad prism (Version 9.0.0). Continuos analysis was completed by Oridinary one-way ANOVA where differences between groups were quantified and a p-value greater than 0.05 was considered significant.

## RESULTS

### Identification of individuals and characterization of clinical features

Five unrelated individuals were included in this study. Individual 1 was enrolled in the UDN, subsequently individuals 2 to 5 were identified in unrelated families through GeneMatcher^21^. We have also conducted a search in a large clinical exome/genome database at Baylor Genetics for candidate *RBBP5* variants, which identified two likely benign de novo variants. The five individuals presented with neurodevelopmental features including developmental delay, intellectual disability, and microcephaly. In addition, short stature, musculoskeletal abnormalities, and dysmorphic facies were the common phenotypes observed in this cohort of individuals (**Figure 1**). Four out of five patients were reported to have short stature and microcephaly. Intellectual disability and global developmental delay were seen in all individuals except the youngest. Sensorineural hearing loss and seizure were reported in two different individuals. Abnormalities in fingers and toes were observed in all five individuals. All individuals presented with dysmorphic facies, but these dysmorphic features did not exhibit as a recognizable pattern. Clinical features of the individuals are summarized in **Table 1**. In addition to the five individuals, we identified two additional cases with de novo *RBBP5* variants. These two cases were considered as likely benign because of inconsistent phenotype and benign in silico analysis (**Supplementary table 1 and 2**).

**Figure 1.**
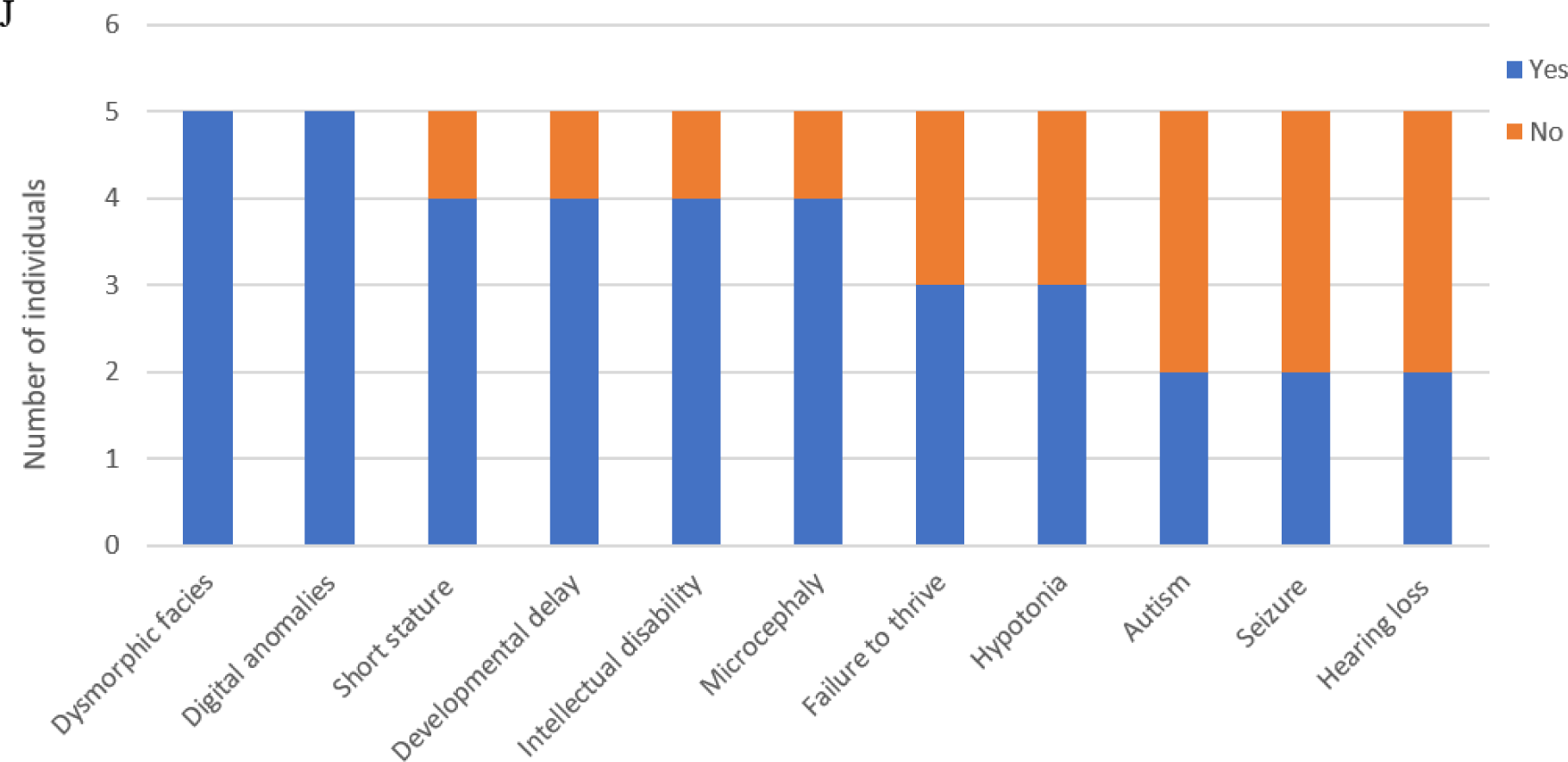
Human subjects with *RBBP5 de novo* variants exhibit a range of clinical features. Dysmorphic features in individual 1 including hypertelorism, high arched eyebrow, long eyelashes, synophrys, and board nasal tip as shown in A; retrognathia, large ear, and a preauricular ear tag as shown in B; bilateral 5^th^ finger clinodactyly and prominent fingertip pad in C. Dysmorphic features in individual 3 as shown in D with midface hypoplasia and cupped ears, clinodacyly in E, and supernumerary teeth in F. Dysmorphic features in individual 4 as shown in G with short and upslanting palpebral fissures, high forehead, anteverted nostrils, and sparse eyebrows. H and I shows the dysmorphic facial features including sparse eyebrows, short nose, long philtrum, small and squared ears, small mouth with thin lips, in individual 5. Common phenotypes are illustrated in J.

**Table 1.**
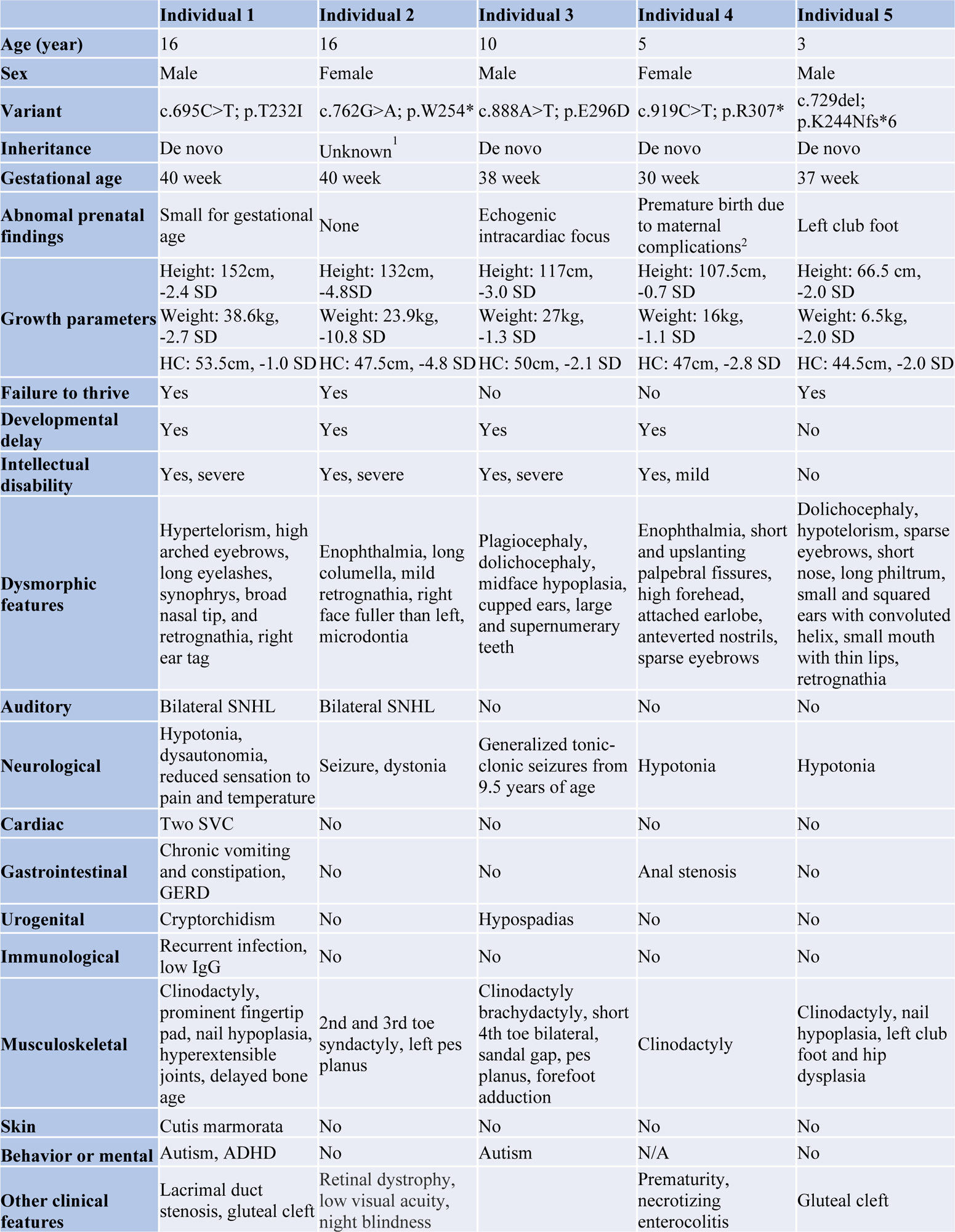
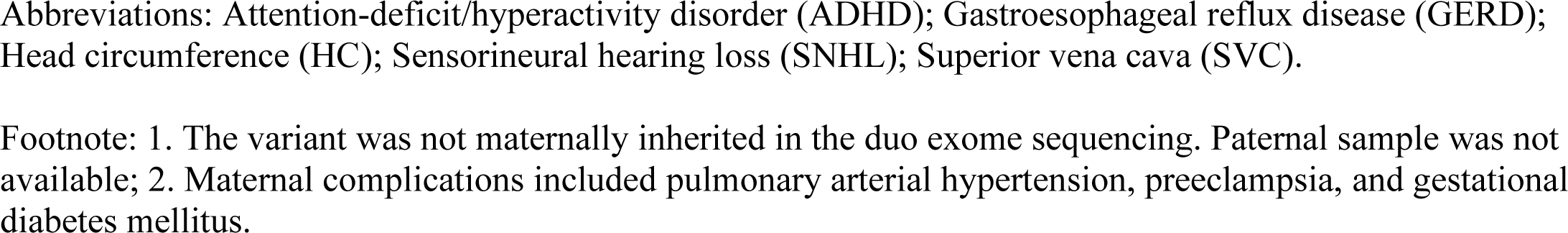
Summary of clinical features in affected individuals.

|  | Individual 1 | Individual 2 | Individual 3 | Individual 4 | Individual 5 |
| --- | --- | --- | --- | --- | --- |
| **Age (year)** | 16 | 16 | 10 | 5 | 3 |
| **Sex** | Male | Female | Male | Female | Male |
| **Variant** | c.695C>T; p.T232I | c.762G>A; p.W254* | c.888A>T; p.E296D | c.919C>T; p.R307* | c.729del;<br>p.K244Nfs*6 |
| **Inheritance** | De novo | Unknown<sup>1</sup> | De novo | De novo | De novo |
| **Gestational age** | 40 week | 40 week | 38 week | 30 week | 37 week |
| **Abnormal prenatal findings** | Small for gestational age | None | Echogenic intracardiac focus | Premature birth due to maternal complications<sup>2</sup> | Left club foot |
| **Growth parameters** | Height: 152cm, -2.4 SD | Height: 132cm, -4.8SD | Height: 117cm, -3.0 SD | Height: 107.5cm, -0.7 SD | Height: 66.5 cm, -2.0 SD |
|  | Weight: 38.6kg, -2.7 SD | Weight: 23.9kg, -10.8 SD | Weight: 27kg, -1.3 SD | Weight: 16kg, -1.1 SD | Weight: 6.5kg, -2.0 SD |
|  | HC: 53.5cm, -1.0 SD | HC: 47.5cm, -4.8 SD | HC: 50cm, -2.1 SD | HC: 47cm, -2.8 SD | HC: 44.5cm, -2.0 SD |
| **Failure to thrive** | Yes | Yes | No | No | Yes |
| **Developmental delay** | Yes | Yes | Yes | Yes | No |
| **Intellectual disability** | Yes, severe | Yes, severe | Yes, severe | Yes, mild | No |
| **Dysmorphic features** | Hypertelorism, high arched eyebrows, long eyelashes, synophrys, broad nasal tip, and retrognathia, right ear tag | Enophthalmia, long columella, mild retrognathia, right face fuller than left, microdontia | Plagiocephaly, dolichocephaly, midface hypoplasia, cupped ears, large and supernumerary teeth | Enophthalmia, short and upslanting palpebral fissures, high forehead, attached earlobe, anteverted nostrils, sparse eyebrows | Dolichocephaly, hypotelorism, sparse eyebrows, short nose, long philtrum, small and squared ears with convoluted helix, small mouth with thin lips, retrognathia |
| **Auditory** | Bilateral SNHL | Bilateral SNHL | No | No | No |
| **Neurological** | Hypotonia, dysautonomia, reduced sensation to pain and temperature | Seizure, dystonia | Generalized tonic-clonic seizures from 9.5 years of age | Hypotonia | Hypotonia |
| **Cardiac** | Two SVC | No | No | No | No |
| **Gastrointestinal** | Chronic vomiting and constipation, GERD | No | No | Anal stenosis | No |
| **Urogenital** | Cryptorchidism | No | Hypospadias | No | No |
| **Immunological** | Recurrent infection, low IgG | No | No | No | No |
| **Musculoskeletal** | Clinodactyly, prominent fingertip pad, nail hypoplasia, hyperextensible joints, delayed bone age | 2nd and 3rd toe syndactyly, left pes planus | Clinodactyly brachydactyly, short 4th toe bilateral, sandal gap, pes planus, forefoot adduction | Clinodactyly | Clinodactyly, nail hypoplasia, left club foot and hip dysplasia |
| **Skin** | Cutis marmorata | No | No | No | No |
| **Behavior or mental** | Autism, ADHD | No | Autism | N/A | No |
| **Other clinical features** | Lacrimal duct stenosis, gluteal cleft | Retinal dystrophy, low visual acuity, night blindness |  | Prematurity, necrotizing enterocolitis | Gluteal cleft |

Abbreviations: Attention-deficit/hyperactivity disorder (ADHD); Gastroesophageal reflux disease (GERD); Head circumference (HC); Sensorineural hearing loss (SNHL); Superior vena cava (SVC).
Footnote: 1. The variant was not maternally inherited in the duo exome sequencing. Paternal sample was not available; 2. Maternal complications included pulmonary arterial hypertension, preeclampsia, and gestational diabetes mellitus.

### Identification and analysis of variants

All five *de novo* heterozygous variants in *RBBP5* (NM_005057.4) were identified through trio whole exome or genome sequencing, including two missense variants p.T232I and p.E296D and three truncating variants p.W254*, p.K244Nfs*6 and p.R307*. *RBBP5* has a pLI score of 1 and missense Z score of 4.64 ^22,23^, suggesting intolerance to truncating and missense changes respectively. In addition, all five variants had not been observed in the gnomAD database^22^. *In silico* variant analysis results were inconsistent in their pathogenicity predictions for the missense variants (Supplementary table 2). Interestingly, all five variants are located in a small region between WD40 repeat domains 4 and 6 in *RBBP5.* This region is predicted to be intolerant to changes based on MetaDome analysis, indicating a potential hotspot for pathogenic variants (**Figure 2A**)^24^. The missense variants affect amino acids that are well-conserved across species from human to *Drosophila* (**Figure 2B**). Expression of the RBBP5 protein with missense variants in HEK293T cells resulted in the full length protein expressed at a similar level compared to that with wild type construct based on Western blotting (**Figure 2C**). However, the single nucleotide deletion at nucleotide 729 in *RBBP5* produced a frameshift product that terminates prematurely and showed no detectable protein expression. Similarly, the C>T change at nucleotide 919 led to premature termination without protein expression (**Figure 2C**). In structural analysis, we found the threonine 232 residue is located in one of the WD40 repeats that forms the interface with the ubiquitin-conjugated histone 2B lysine-120 (H2BK120), and T232 in RBBP5 is predicted to be the key residue mediating the interaction between RBBP5^WD40^ and the ubiquitin-conjugated H2BK120 (**Figure 2D and E**). Similarly, the E296 residue is also located at a crucial position mediating intereaction of RBBP5^WD40^ and histone 2B (**Figure 2F**).

**Figure 2.**
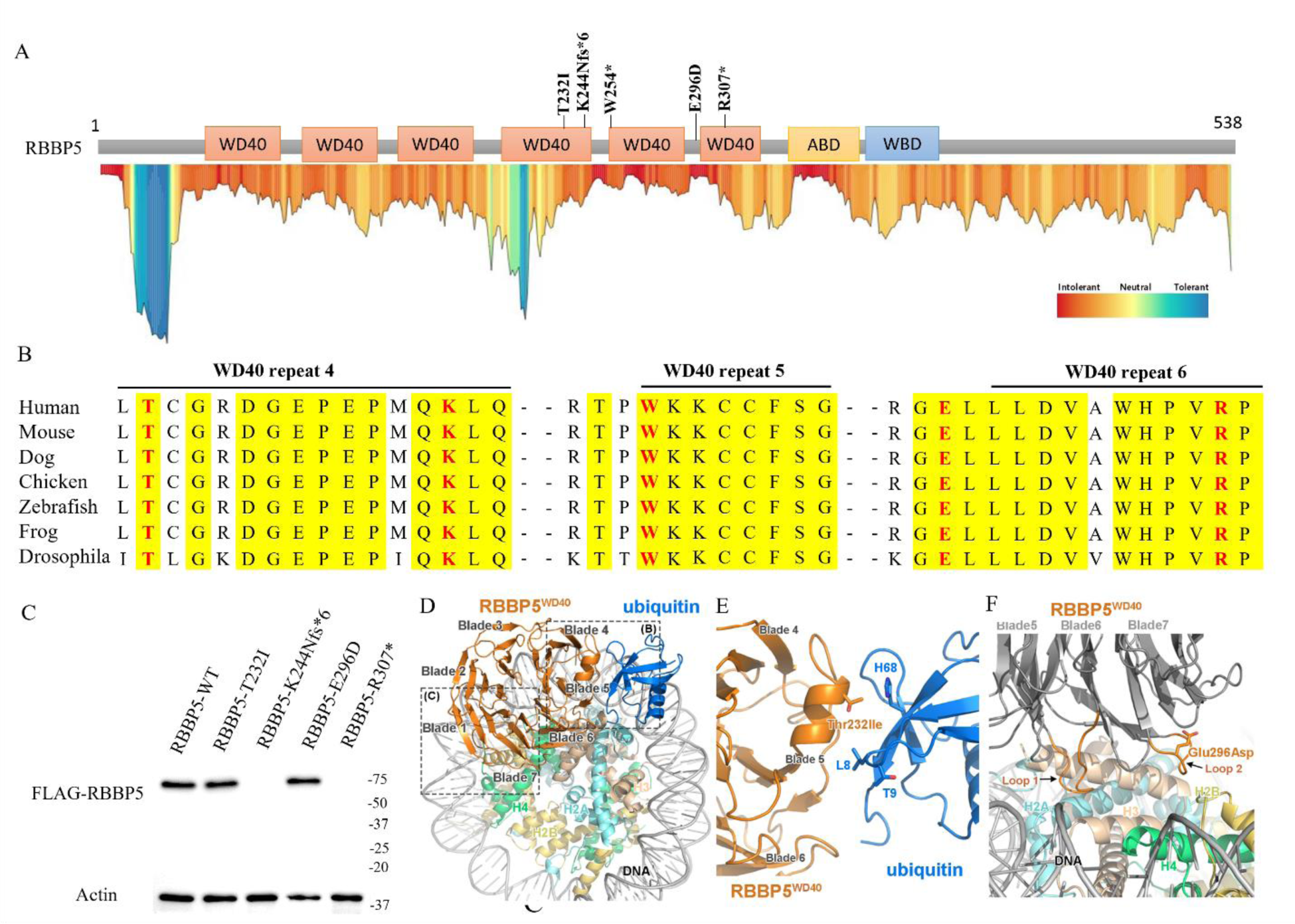
Bioinformatic and structual variant analysis. The functional domains of RBBP5 and position of variants in MetaDome are shown in A. The evolutionarily conserved residues affected by variants are shown in B. The protein expression of FLAG-tagged human RBBP5 reference and variants were shown in C. Structure analysis of T232 and E296 were performed in RBBP5^WD40^ of the cryo-EM structure of MLL3-ubNCP complex. The overall structure of RBBP5^WD40^ complexed with an nucleosome core particle mono-ubiquitinated at the Lys 120 of histone H2B (ubNCP). The RBBP5 WD40 repeat 4 is sandwiched between ubiquitin and core histones. The RBBP5^WD40^ is shown in orange and ubiquitin in blue (D). Detailed view of the recognition interface of RBBP5^WD40^-ubiquitin. Residue Thr232Ile, which is located on the α-helix-containing loop of RBBP5^WD40^ blade 5, lies close to residues Leu8, Thr9 and His68 of ubiquitin (E). All these residues are shown in stick model. Detailed view of the interaction interface between RBBP5^WD40^ and histone H2B-H4. Two loops (Loop 1 and Loop 2), which connect the WD40 propeller blades 5, 6 and 7, interact with nucleosome directly. Residue Glu296Asp is located on Loop 2 and shown in stick model (F).

### Overexpression of human *RBBP5* missense variants in *Drosophila* results in microcephaly

For this study, we performed overexpression and rescue experiments with the human *RBBP5* cDNA to determine if the variants have a functional consequence *in vivo.* The human *RBBP5* is orthologous to *Rbbp5* in *Drosophila* (DIOPT score: 14/16)^25^. In flies, *Rbbp5* is a member of the trithorax complex which is required for differentiation of neural lineages^26,27^. Importantly, neuronal fate determination is similar between humans and flies^28^. Previous research has reported complete loss of H3K4 trimethylation (H3K4me3) in *Rbbp5* mutant clones^27^. The resulting *Rbbp5* loss-of-function phenotype includes an inability to maintain type II neuroblast identity that gives rise to intermediate progenitor cells. *Rbbp5* null neuroblasts instead express markers for type I neuroblasts^27^. Intermediate progenitor cells undergo several rounds of self-renewal before being committed to a ganglion mother cell that will terminally differentiate into two neurons. Critically, type I neuroblasts are not capable of generating intermediate progenitor cells. This study also demonstrated that neuronal identity and H3K4me3 levels can be restored by the expression of the full length Rbbp5^27^. A shift from type II to type I neuroblast identity could result in a decrease in the overall number of neurons inducing a microcephaly phenotype in flies similar to the phenotype of the individuals included in this study.

We first determined whether overexpression of the human reference *RBBP5* cDNA (NM_005057.4) (*RBBP5^Ref^*), or variants p.T232I (*RBBP5^T232I^*), p.E296D (*RBBP5^E296D^*) by ubiquitous or tissue specific *GAL4* drivers causes phenotypes in the fly. In the *GAL4-UAS* system, a *GAL4* transcriptional activator protein is fused to the promoter of a gene of interest. When paired with a construct containing an upstream activation sequence (UAS), this drives expression of the construct based on the spatial and temporal expression pattern of the gene^29^. In human protein overexpression paradigms, the endogenous *Drosophila* gene is unaffected thus enabling the detection of gain- or loss-of-function mechanisms. Gain-of-function phenotypes would appear more severe than wild-type and loss-of-function would appear wild-type since the fly gene is expressed in the genetic background^30^.

We overexpressed *RBBP5^Ref^*, *RBBP5^T232I^,* or *RBBP5^E296D^* using *Actin*-(*Act-*, strong ubiquitous), *daughterless-* (*da-,* weak ubiquitous) *eyeless-* (*ey-,* developing visual system and parts of the head and brain)*, elav-* (neuron), and *repo-* (glia) *GAL4* lines for tissue specific expression. We then scored lethality by developmental stage where the *Drosophila* life cycle is ten days long at 25°C. Overexpression of *RBBP5^Ref^* or either missense variant with *Actin^GAL4^*is lethal in the third instar larval (L3) stage (**Figure 3A**). Flies undergo embryo, larval, pupal, and adult stages and larval development is marked by progression through L1-L3 stages, with L3 spanning three days before pupation^31^. Expression of *RBBP5^Ref^* with *da^GAL4^* is again L3 lethal, however expression of *RBBP5^T232I^* or *RBBP5^E296D^* is pupal lethal (**Figure 3A**). We observe no decrease in viability with *ey-, elav-,* and *repo-GAL4* drivers (**Figure 3A**). Both *Actin* and *daughterless* are expressed early in development, and these data support previous findings that *Rbbp5* has a critical role early in development. Disruption later in development with *elav-* or *repo^GAL4^* induces no phenotype because critical functions have already been carried out when neuronal and glial specific genes are expressed. Importantly, since the fly *Rbbp5* is being expressed in the genetic background in these human cDNA overexpression experiments, observing a loss of lethality that is induced by overexpression of the *RBBP5^Ref^* suggests both *RBBP5^T232I^* and *RBBP5^E296D^* are loss-of-function alleles.

**Figure 3.**
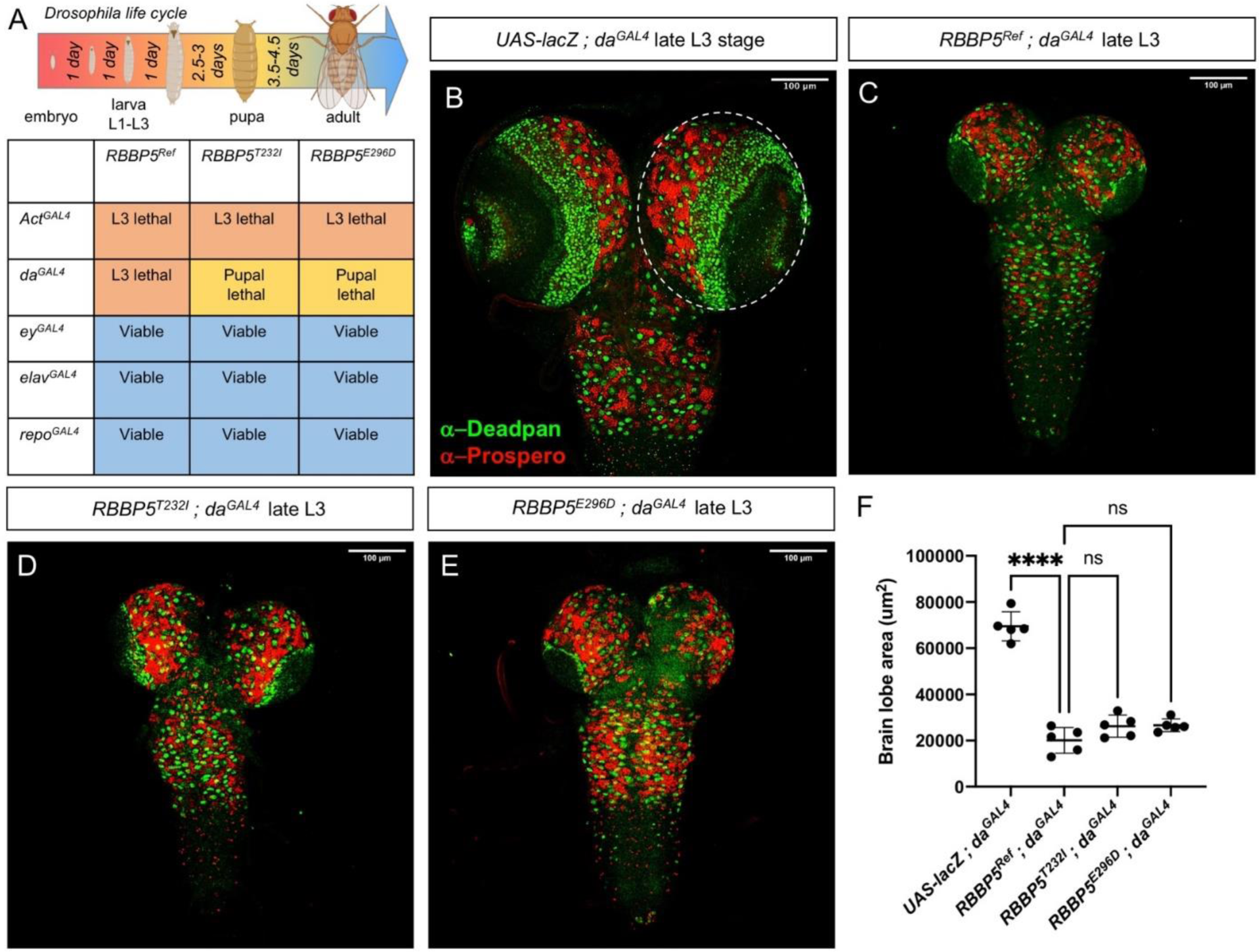
Ubiquitous expression of *RBBP5* in flies induces lethality and results in microcephaly in the larval developmental stage. The *Drosophila* life cycle is approximately 10 days long at 25°C. Expression of the human *RBBP5* or either missense variant with a strong ubiquitous driver (*Actin^GAL4^*) is larval lethal (*Actin^GAL4^ / RBBP5^Ref^* observed in 0/128 F1 progeny, o/e=0.0; *Actin^GAL4^ / RBBP5^T232I^* observed in 0/144 F1 progeny, o/e=0.0; *Actin^GAL4^ / RBBP5^E296D^* observed in 0/277, o/e=0.0). With a weak ubiquitous driver (*da^GAL4^*), expression of the human reference is larval lethal, but expression of p.T232I or p.E296D is pupal lethal. Expression with tissue specific (*ey-*, *elav-*, or *repo^GAL4^*) drivers does not affect viability in A. Representative developmentally staged control late L3 brains (*UAS-lacZ; da^GAL4^*) with Deadpan staining of neuroblasts and intermediate progenitor cells in green and Prospero staining of neural progenitors in red in B. Experimental *RBBP5* reference (*RBBP5^Ref^; da^GAL4^*) brain shown in C, *RBBP5^T232I^* (*RBBP5^T232I^; da^GAL4^*) in D and *RBBP5^E296D^* (*RBBP5^E296D^; da^GAL4^*) in E. Quantification of ubiquitous overexpression (*da^GAL4^*) of *RBBP5^Ref^*, *RBBP5^T232I^*, and *RBBP5^E296D^* compared to *UAS-lacZ* (one-way ANOVA, ns, p>0.05, *p<0.05, **p<0.01, ***p<0.001, ****p<0.0001) in F. Created with Biorender.com.

We investigated brain development during the larval stage because *daughterless* overexpression of *RBBP5^Ref^* is L3 lethal while expression of *RBBP5^T232I^* or *RBBP5^E296D^* is pupal lethal. We dissected L3 brains using *da^GAL4^* to express a neutral cDNA (*da^GAL4^; UAS-lacZ*) as a control and immunostained for markers of progenitor lineages, Deadpan (neuroblasts and intermediate progenitor cells) and Prospero (differentiating neurons)^27^ (**Figure 3B**). Deadpan-positive intermediate progenitor cells are present in the optic lobes of control larvae, however when *RBBP5^Ref^* is expressed, L3 stage matched brains are severely reduced in size with a loss of intermediate progenitor cells in the optic lobes of the brain (**Figure 3C, 3F**). Interestingly, while overexpression of either missense variant is pupal lethal, *RBBP5^T232I^* and *RBBP5^E296D^* L3 brains also fail to develop to control lobe size and are not significantly larger than *RBBP5^Ref^* brains (**Figure 3D-F**) despite the observed difference in lethality staging. These results suggest that growth phenotypes may not be directly linked, and overall growth and brain development could be directed by different pathways including RBBP5/Rbbp5.

We therefore dissected L3 brains using the strong ubiquitous driver *Actin^GAL4^* which induces L3 lethality with *RBBP5^Ref^, RBBP5^T232I^,* or *RBBP5^E296D^.* We again performed immunostainings to label neural progenitor lineages with Deadpan and Prospero antibodies^32^. Early and late L3 control (*Actin^GAL4^; UAS-lacZ*) larvae were used to identify neural development occurring throughout the L3 stage (**Supplementary Figure 1A-B**). Late L3 brains that express *RBBP5^Ref^* are less developed in size compared to control stage-matched wandering L3 larva (**Supplementary Figure 1C, F**). Late L3 brains that express *RBBP5^T232I^* or *RBBP5^E296D^*also display a small brain (**Supplementary Figure 1D-F**). Ubiquitous expression of *RBBP5* leads to dramatic reduction in brain size with no significant differences in size between *RBBP5^Ref^*, *RBBP5^T232I^*, or *RBBP5^E296D^* (**Supplementary Figure 1F**). Overall, these data suggest that expression of the human RBBP5 interrupts function of the fly Rbbp5 protein and results in a strong microcephaly phenotype in the fly.

We also observed changes in overall larval size and development upon ubiquitous overexpression of *RBBP5*. Larval developmental progression is a highly stereotyped pattern and a failure to reach developmental stages indicates possible dysregulation of factors that direct development^33^. Late L3 larvae ubiquitously expressing *RBBP5^Ref^, RBBP5^T232I^,* or *RBBP5^E296D^*with *Actin^GAL4^* exhibit reduced overall body size compared to control larvae expressing *UAS-lacZ* (**Supplementary Figure 2A**). Control wandering L3 (**Supplementary Figure 2B**) develop mature anterior and posterior spiracles indicating that the late L3 stage has been reached^34^.

Severe growth phenotypes are seen upon expression of the human reference cDNA. Expression of either *RBBP5^Ref^* (**Supplementary Figure 2C**) or *RBBP5^T232I^* (**Supplementary Figure 2D**) results mature posterior spiracle formation but failure of the posterior spiracles to develop. In *RBBP5^E296D^* expressing larvae however, anterior and posterior spiracles successfully develop similar to controls (**Supplementary Figure 2E**). These results indicate an inability of p.E296D to induce the developmental phenotype seen in reference expressing larvae suggesting that in the context of larval developmental progression, p.E296D could be a strong loss-of-function allele. To investigate the effect of our *RBBP5* variants on trimethylation in the L3 developmental stage, we confirmed RBBP5 expression with *Actin^GAL4^* and quantified H3K4me3 compared to total H3 (**Supplementary Figure 2F-G**). A significant reduction in H3K4me3 is observed in *RBBP5^Ref^, RBBP5^T232I^,* and *RBBP5^E296D^* compared to *UAS-lacZ* control larvae (**Supplementary Figure 2G**). These results suggest that dysregulation of Rbbp5 results in the failure to express developmental genes that are critical for the progression through the L3 developmental stage. These data confirm that the human RBBP5 can interact with the fly trithorax complex members and that expression of human alleles can impact H3K4me3 levels in the fly.

### Tissue-specific *RBBP5* expression in *Drosophila* results in a small eye phenotype

To assess *RBBP5* function in the eye we drove the variants using the *eyeless^GAL4^* and quantified eye size in the adult stage. The eyes of *RBBP5^Ref^* or *RBBP5^E296D^* flies are smaller than *UAS-lacZ* or *RBBP5^T232I^* (**Figure 4A-E**). These data suggest that in the eye there is a toxic effect of overexpression of *RBBP5^Ref^* and *RBBP5^E296D^* which is not as severe in *RBBP5^T232I^* suggesting that in the context of eye development, p.T232I could be a strong loss-of-function allele. Overall, these experiments conclude that overexpression of the human *RBBP5* is toxic and results in growth phenotypes including microcephaly in the brain and reduced size in the body and eye.

**Figure 4.**
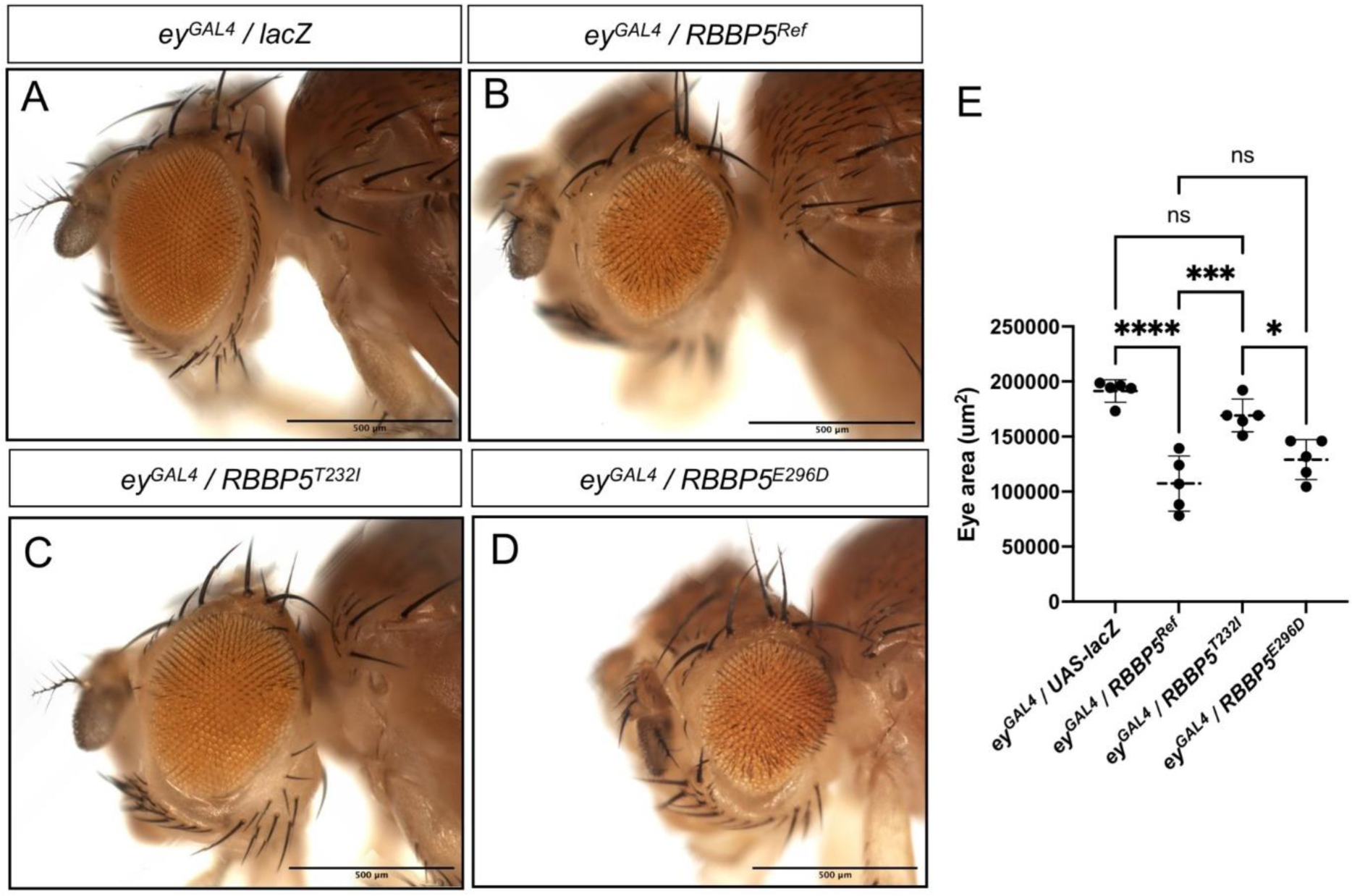
Human *RBBP5* expression induces a small eye phenotype not recapitulated by the p.T232I variant. Overexpression with *eyeless-GAL4* (*ey^GAL4^*) in a control line (*ey^GAL4^ / UAS-lacZ*) as shown in A, *ey^GAL4^ / RBBP5^Ref^* in B, *ey^GAL4^ / RBBP5^T232I^* in C, and *ey^GAL4^ / RBBP5^E296D^* in D. Expression of *RBBP5^Ref^* and *RBBP5^E296D^* results in a small eye phenotype compared to *UAS-lacZ*. Expression of *RBBP5^T232I^*does not induce a small eye phenotype and eye size is not significantly different than controls (one-way ANOVA, ns p>0.05, *p<0.05, **p<0.01, ***p<0.001, ****p<0.0001) in E.

### Overexpression of *Rbbp5* leads to wing patterning defects

Next, we compared human and fly cDNA overexpression. We generated the orthologous fly *Rbbp5* constructs and created the p.T232I and p.E296D homologous variants, p.T231I (*Rbbp5^T231I^*) and p.E295D (*Rbbp5^E295D^*) respectively. We compared this to an HA-tagged version of *RBBP5* (*RBBP5-HA*). Ubiquitous expression of *Rbbp5, Rbbp5^T231I^,* or *Rbbp5^E295D^*with *Actin^GAL4^* or *da^GAL4^* is viable, but *Actin^GAL4^* induces ectopic wing vein formation that is not fully penetrant at 25°C (**Supplementary Figure 3A**). Since the GAL4-UAS system is temperature dependent, we increased the culture temperature, and upon expression of *Rbbp5* ectopic wing vein formation is fully penetrant at 29°C, but the phenotypes of *Rbbp5^T231I^* or *Rbbp5^E295D^* are not fully penetrant (**Supplementary Figure 3B-E**). Expression of *RBBP5-HA* is pupal lethal with *Actin^GAL4^* and viable with *da^GAL4^*but ectopic wing vein formation is present (**Supplementary Figure 3A**). Ectopic wing vein formation is observed with both *da^GAL4^* and *Actin^GAL4^* compared to the laboratory control strain Canton S, however *Actin^GAL4^* escaper flies are rarely observed (**Supplementary Figure 3F-H**). The ectopic wing vein formation seen upon overexpression of *RBBP5-HA* is similar to the phenotypes observed with *Rbbp5* suggesting a disruption of factors that direct wing development when the fly or human cDNA is overexpressed. Furthermore, expression of the wild-type fly cDNA induces fully penetrant wing patterning defects while either missense variant p.T231I or p.E295D results in wing phenotypes that are not fully penetrant again indicating a loss-of-function mechanism in the context of the fly variants.

### Expression pattern of *Rbbp5* in the fly brain

To determine the *Rbbp5* expression pattern and further explore the function of the variants, we replaced the open reading frame of *Rbbp5* with the *KozakGAL4* sequence (*Kozak sequence-GAL4-polyA-FRT-3XP3EGFP-polyA-FRT*)^28^. This effectively removes *Rbbp5* and leads to GAL4 expression in a similar spatial and temporal expression pattern (hereafter *Rbbp5^Kozak^ ^GAL4^*). The GAL4 transcriptional activator protein will bind UAS containing constructs to drive expression of a reporter protein or human cDNA to determine if the human protein can rescue loss of the *Drosophila* protein and whether that ability is impaired by the variant^29^. We used the *Rbbp5^Kozak^ ^GAL4^*allele to determine the expression pattern of *Rbbp5* in the developing nervous system. We crossed the *Rbbp5^Kozak^ ^GAL4^* to a *UAS-mCherry NLS* reporter line and determined that *Rbbp5* is expressed in a subset of Elav*-*positive neurons (**Figure 5A-C**) and Repo-positive glia (**Figure 5D-F**) in the optic lobes and ventral nerve cord in the larval brain. Confirming *Rbbp5* expression in both neurons and glia supports the canonical function of *Rbbp5* to direct neuronal fate because type II neuroblasts give rise to both neurons and glia.

**Figure 5.**
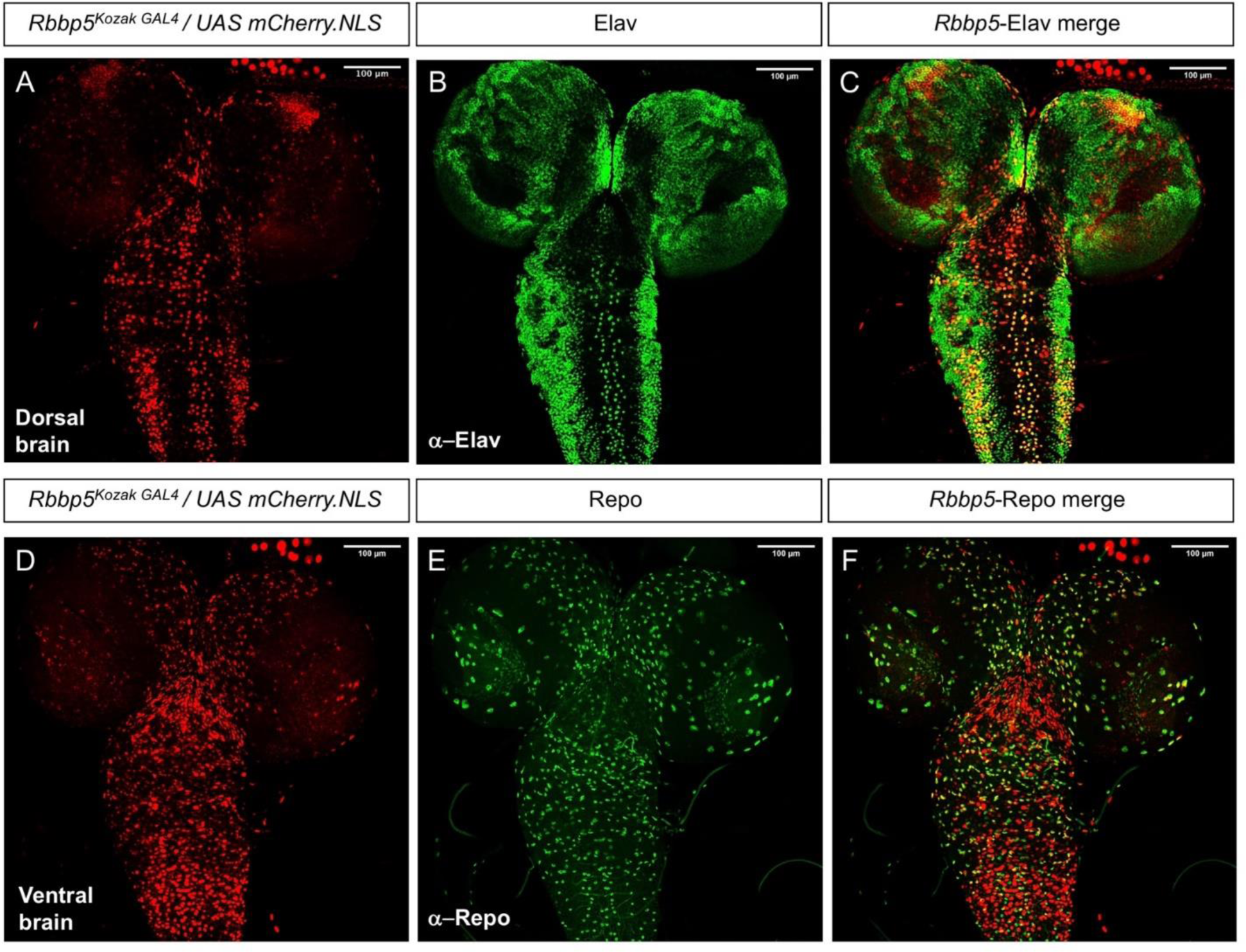
*Rbbp5* is expressed in a subset of neurons and glia in the *Drosophila* brain. *Rbbp5^Kozak^ ^GAL4^ / UAS mCherry.NLS* expression pattern in the dorsal larval brain shown in A. Elav expression in B, and merge in C with co-localization in the ventral nerve cord and optic lobes of the central brain. *Rbbp5^Kozak^ ^GAL4^ / UAS mCherry.NLS* expression pattern in the ventral larval brain shown in D. Repo expression in E, and merge in F with co-localization in the ventral nerve cord and optic lobes.

### *RBBP5* missense variants produce less severe phenotype in microcephaly rescue experiment

Next, we assessed the phenoptypes associated with loss of *Rbbp5* function. We dissected L3 brains of *Rbbp5^Kozak^ ^GAL4^* heterozygous mutant animals and a laboratory control strain (*y^1^ w\**) and again immunostained for markers of progenitor lineages, Deadpan and Prospero^30^. Deadpan-positive intermediate progenitor cells are present in the optic lobes *Rbbp5^Kozak^ ^GAL4^* heterozygous animals (**Figure 6A**) and there is no difference in brain size between *Rbbp5^Kozak^ ^GAL4^* / + and *y^1^ w\** controls (**Figure 6F**). However, in *Rbbp5* null animals (*Rbbp5^Kozak^ ^GAL4^ / Df(3L)BSC447*) (*Df(3L)BSC447* is a 125 kB deficiency that encompasses the *Rbbp5* locus), loss of *Rbbp5* is pupal lethal. Furthermore, the development of intermediate progenitor cell is severely impaired, and brain size is reduced in the L3 developmental stage (**Figure 6B, F**). When we attempt to rescue this microcephaly phenotype with *Rbbp5^Kozak^ ^GAL4^* driven expression of *RBBP5^Ref^*, it fails to rescue the loss of the fly *Rbbp5*, moreover there is a further reduction in brain size and more severely impacted intermediate progenitor cell population (**Figure 6C, F**). Therefore the *RBBP5^Ref^* transgene not only fails to rescue, but also exacerbates the loss-of-function phenotypes.

**Figure 6.**
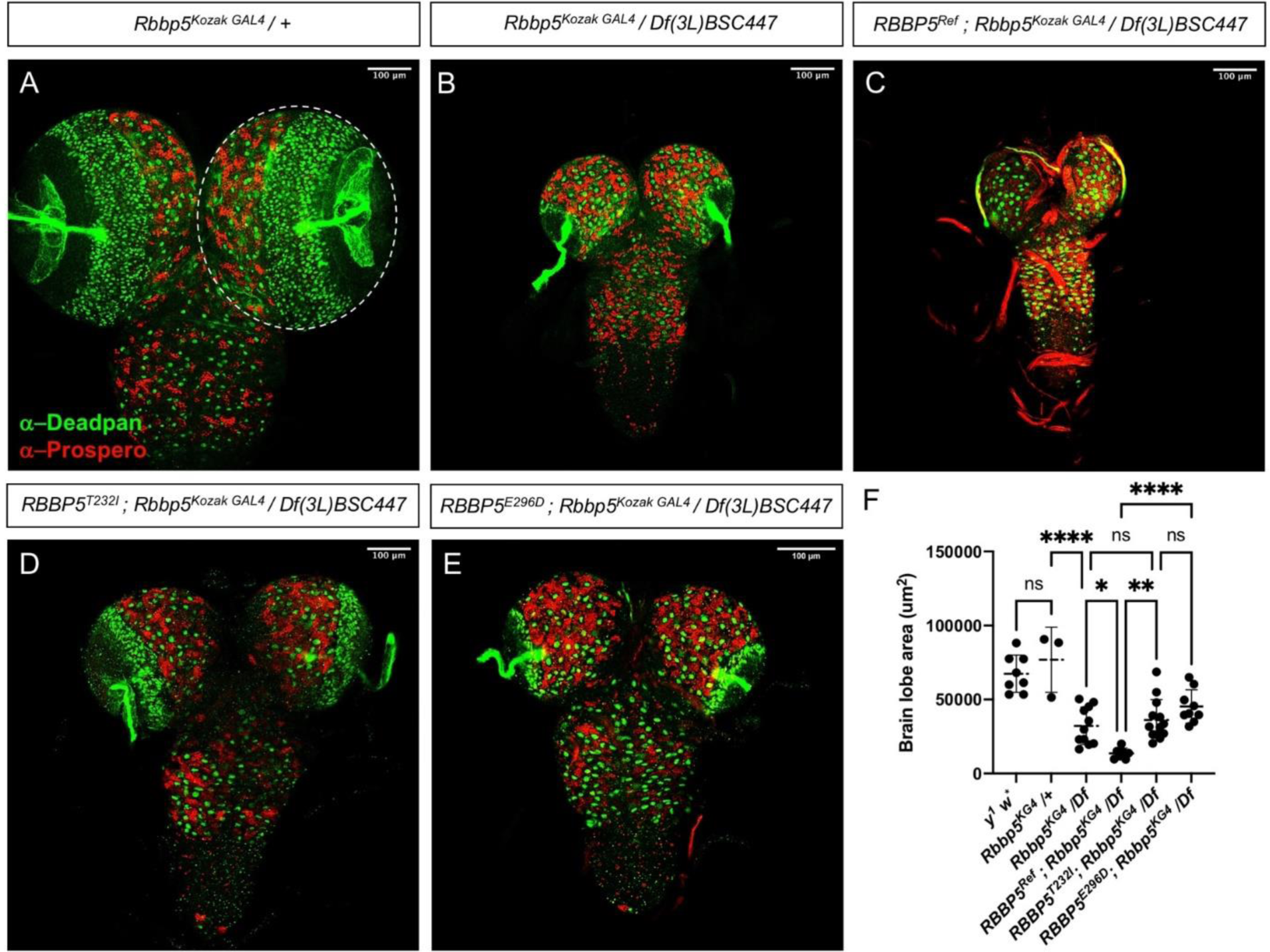
*RBBP5* human transgenes fail to rescue loss of *Drosophila Rbbp5*. Heterozygous L3 *Rbbp5* loss-of-function controls (*Rbbp5^Kozak^ ^GAL4^ / +*) with Deadpan positive neuroblasts and intermediate progenitor cells in green and Prospero positive neural progenitor cells in red in A. Homozygous loss-of-function with *Rbbp5^Kozak^ ^GAL4^* crossed to a deficiency line *Df(3L)BSC447* that includes the *Rbbp5* locus (*Rbbp5^Kozak^ ^GAL4^ / Df(3L)BSC447*) in B. Attempted rescue with the human *RBBP5^Ref^* (*RBBP5^Ref^; Rbbp5^Kozak^ ^GAL4^/ Df (3L)BSC447*) in C, *RBBP5^T232I^*(*RBBP5^T232I^; Rbbp5^Koza^ ^kGAL4^/ Df(3L)BSC447*) in D, *RBBP5^E296D^* (*RBBP5^E296D^; Rbbp5^Kozak^ ^GAL4^/Df(3L)BSC447*) in E. Quantification of the microcephaly phenotype (brain area) by genotype (one-way ANOVA, ns p>0.05, *p<0.05, **p<0.01, ***p<0.001, ****p<0.0001) in F. *RBBP5^Ref^*, *RBBP5^T232I^*, and *RBBP5^E296D^* fail to rescue loss of the *Drosophila Rbbp5. RBBP5^Ref^* expression induces a significantly more severe microcephaly phenotype than *Rbbp5^Kozak^ ^GAL4^/ Df (3L)BSC447*, and brain size of *Rbbp5^T232I^; Rbbp5^KozakGAL4^/ Df(3L)BSC447* or *Rbbp5^E296D^; Rbbp5^KozakGAL4^/ Df(3L)BSC447* larvae is not significantly different than *Rbbp5^KozakGAL4^/ Df (3L)BSC447*.

Next we attempted the rescue experiment with *RBBP5^T232I^* (**Figure 6D**) or *RBBP5^E296D^* (**Figure 6E**) and these transgenes also do not rescue, but do not produce the more severe phenotypes observed with *RBBP5^Ref^***(Figure 6C**). These results indicate that p.T2321 or p.E296D cannot rescue loss of *Rbbp5* but impair development less severely than the reference cDNA (**Figure 6F**). This again indicates that overexpression of the *RBBP5^Ref^* is toxic and that *RBBP5^T232I^ and RBBP5^E296D^* are less toxic even when expressed under the control of endogenous *Rbbp5* promoter. In summary, total brain lobe area was reduced in the *Rbbp5* mutants and was not rescued by the human reference nor the p.T232I or p.E296D transgenes. However, *RBBP5^Ref^* produces even stronger lobe size reduction than *Rbbp5* null larvae. Furthermore, there is no significant difference in brain lobe area between either *RBBP5^T232I^* or *RBBP5^E296D^* in a null genetic background and *RBBP5* null lobe size, again supporting the finding that p.T232I and p.E296D are loss-of-function alleles (**Figure 6F**). This is consistent with the phenotypes observed in overexpression experiments with *RBBP5^T232I^* and *RBBP5^E296D^* failing to induce the toxic effects of *RBBP5^Ref^*. Ubiquitous expression of *RBBP5^Ref^* induces earlier lethality than expression of *RBBP5^T232I^* or *RBBP5^E296D^*, and impaired growth phenotypes are present upon expression of the human *RBBP5* reference that are not present when the missense variants are expressed indicating a loss-of-function mechanism for the p.T232I and p.E296D variants.

## Discussion

In this study, we identified five affected individuals with *de novo* heterozygous variants in one of the COMPASS core members *RBBP5*. We propose that haploinsufficient loss of *RBBP5* is responsible for the neurodevelopmental disorder presented here. This is consistent with the fact that *RBBP5* has a pLI score of 1, and other disorders of epigenetic machinery classically are due to haploinsufficiency^5^. In fact, the p.T232I and p.E296D missense variants met the criteria for a pathogenic variant based on two strong criteria from the American College of Medical Genetics and Genomics variant interpretation guideline (PS2 confirmed *de novo* and PS3 *in vitro* or *in vivo* functional studies supportive of a damaging effect)^35^. In addition, the absence of these five variants in the population database and the low rate of benign missense variation in *RBBP5* are also supportive for the pathogenic variant classification.

The COMPASS protein complex consists of four core members: RBBP5, WDR5, ASH2L, and one of the six methyltransferases^36^. The methyltransferases have the enzymatic SET1 domain to methylate H3K4, while RBBP5 functions to modulate the activity of the complex and mediate the interaction between the nucleosome and the complex^4,37^. Our structural analysis showed both the T232 and E296 residues are located at critical positions of the interface between RBBP5 and the histone H2B, which has been known to be essential for the recruitment of COMPASS to the nucleosome^38–40^. The mutations in T232 and E296 are likely to interrupt the interaction between RBBP5 and H2B, resulting in the dysregulation of downstream target genes.

Kabuki syndrome is one of the most common disorders in the epigenetic machinery^6^. *KMT2D*, one of the methyltransferases in the COMPASS, is the major disease gene for the Kabuki syndrome. Nevertheless, about 20-30% of clinically diagnosed Kabuki syndrome patients have negative genetic testing^6^. It has been hypothesized that a pathogenic variant in other COMPASS members could result in a disorder that phenotypically resembles Kabuki syndrome^7^. We observe some striking similarities in phenotypes between our probands and those typically seen in patients with Kabuki syndrome, such as the neurodevelopmental features, microcephaly, short stature, hypotonia, sensorineural hearing loss, and seizure^41^. However, there are phenotypes such as congenital cardiac defects and the characteristic dysmorphic facial features in Kabuki syndrome that are absent in our probands. Given KMT2D is only one of the six methyltransferases to which RBBP5 binds, it is reasonable to expect differences in the clinical spectrum between *KMT2D* and *RBBP5*-related disorders.

The *Drosophila* model has previously been used to confirm the functional mechanism of variants in SET domain containing methyltransferases^42^. The fly Rbbp5 interacts with trithorax proteins, a coactivator complex that maintains gene activation through H3K4 methylation^27^. The Kabuki syndrome implicated methyltransferase *KMT2D* is homologous to *trithorax related* (*trr*), a gene that is important for eye development and hormone responsive development^43^. In addition to RBBP5, another COMPASS member ASH2L homolog, ash2, also interacts with Trr and the ecdysone receptor (EcR) to direct molting and metamorphosis^44^. Notably, *ash2* mutants also exhibit neural and optic lobe developmental defects^45^.

We present evidence that the *RBBP5* p.T232I and p.E296D variants are hypomorphic loss-of-function alleles. We observed earlier lethality upon ubiquitous expression of the human reference than either missense variant. We also observed variant specific loss-of-function phenotypes using ubiquitous and tissue specific overexpression. We identified that *Rbbp5* is expressed in both neurons and glia in the developing *Drosophila* brain. We found that loss of *Rbbp5* results in microcephaly in the larval stage and confirmed that this loss is lethal. Unfortunately, the human *RBBP5* is unable to rescue loss of the *Drosophila Rbbp5* gene, however we did find variant specific differences when the *RBBP5* transgenes are expressed. A similar microcephaly phenotype is induced upon co-expression of either p.T232I or p.E296D in an *Rbbp5* null background that is not as severe as co-expression of the human reference. Moreover, expression of p.T232I or p.E296D in a null background induces the same microcephaly phenotype as seen in *Rbbp5* null animals confirming that both missense variants are loss-of-function alleles. In addition, expression of the fly and human cDNA can lead to wing patterning defects. Furthermore, expression of the fly p.T231I or p.E295D variants fail to induce fully penetrant wing patterning defects as seen upon overexpression of the fly *Rbbp5.* While we conclude that both missense variants investigated in this study are loss-of-function, the toxicity observed upon overexpression of *RBBP5* suggests *RBBP5* could be a dosage sensitive gene.

We have shown that H3K4 trimethylation is disrupted in *RBBP5* expressing animals, suggesting an inability to activate expression of key developmental genes. We observed variant specific developmental abnormalities in larvae ubiquitously overexpressing *RBBP5* including microcephaly and overall growth phenotypes. Therefore, the p.T232I and p.E296D variants disrupt the function of the COMPASS complex possibly due to substitution of critical residues resulting in an inability to trimethylate H3K4 to direct downstream transcriptional activation. Since we observe unique tissue specific loss-of-function phenotypes between the missense variants, it is possible that downstream gene expression is dysregulated in a variant specific manner. Future transcriptomics studies could identify the critical genes dysregulated by these and additional *RBBP5* variants. Furthermore, inclusion of variants in additional members of the COMPASS complex in transcriptomics studies could begin to identify the target genes responsible for the overlapping and distinct phenotypes involved in the spectrum of observed clinical symptoms.

In summary, we have provided the first evidence for a syndromic neurodevelopmental disorder that is associated with pathogenic variants in *RBBP5*. This study provides a new perspective to the disorders of the epigenetic machinery.

## Data Availability Statement

The authors confirm that the data supporting the findings of this research are available within the manuscript or available upon request.

## Acknowledgements

We thank all participating patients and their family members for supporting this study.

## Author Contributions

Conceptualization: Y.H., K.L.J., M.F.W., J.A.M-A.; Data curation: Y.H., J.A.M-A.; Formal analysis: Y.H., K.L.J., M.F.W., J.A.M-A.; Funding acquisition: J.A.M-A., S.F.N., C.G.P., M.F.W., S.Y., H.J.B., Y.H.; Investigation: Y.H., K.L.J., A.Y-W.H., J.W., S.V.J., O.C., A.R., O.B., M.M., M.I., H.X., J.H., C.M., K.B., V.S., A.Y.M-S., F.R., S.S., C.C., J.A.R; Resources: Y.H., K.L.J., O.K., H.J.B., M.F.W. J.A.M-A.; Supervision: S.Y., H.J.B., C.G.P., S.F.N. M.F.W., J.A.M-A.; Writing--original draft: Y.H., K.L.J.; Writing-review and editing: Y.H., K.L.J., A.Y-W.H., J.W., S.V.J., O.C., A.R., O.B., M.M., M.I., H.X., J.H., C.M., K.B., V.S., A.Y.M-S., F.R., S.S., C.C., S.Y., O.K., H.J.B., J.A.R., C.G.S., S.F.N., M.F.W., J.A.M-A.

## Ethics declaration

Probands were recruited through their local referring physicians and the UDN clinical site (UCLA). Individual 1 (UDN903866) was identified through the Undiagnosed Diseases Network (UDN), and individuals 2-4 were identified through GeneMatcher. Prior to inclusion, informed written consent was obtained from the legal guardians of the individuals included in this study for research and publication according to the standards and practices of the institutional review board and ethics committee at UCLA. Documents and consent forms were standardized according to the requirements of the UDN.

---

**Supplemental Table 1.** Likely bengin RBBP5 de novo variants.

|  | Benign variant 1 | Benign variant 2 |
| --- | --- | --- |
| Age (year) | 1 | 7 |
| Sex | Male | Female |
| Variant | c.1490C>G; p.S497C | c.1589-3T>C |
| Inheritance | De novo | De novo |
| Gestational age | N/A | 40 week |
| Abnormal prenatal findings | None | None |
| Growth parameters | N/A | Height: 118.4 cm, -1.33SD |
|  |  | Weight: 21.2kg, -1.0 SD |
|  |  | HC: 55cm, -0.12 SD |
| Failure to thrive | No | Yes, growth hormone deficiency from hypoplastic pituitary |
| Developmental delay | No | No |
| Intellectual disability | No | No |
| Dysmorphic features | No | Triangular shaped face |
| Auditory | No | No |
| Neurological | Seizure, choreoathetosis | No |
| Cardiac | History of cardiorespiratory arrest | Tiny patent ductus arteriosus |
| Gastrointestinal | No | Celiac disease |
| Urogenital | No | No |
| Immunological | No | History of transient hypogammaglobulinemia of infancy |
| Musculoskeletal | No | Generalized joint hypermobility |
| Skin | No | No |
| Behavior or mental | No | No |
| Other clinical features | Thrombocytosis | Mildly elevated serum CK and mildly low ceruloplasmin; elevated AST |

**Supplemental Table 2.** In silico analysis of *RBBP5* de novo variants.

| Refseq<br>(NM_005057.4) | GRCh38<br>(hg38) | Allele<br>frequency<br>(gnomAD) | Consequence | PolyPhen-2 | REVEL | CADD | SpliceAI |
| --- | --- | --- | --- | --- | --- | --- | --- |
| c.695C>T;<br>p.T232I | 1:205100209 | 0 | missense | Benign | 0.256 | 22.9 |  |
| c.729del;<br>p.K244Nfs*6 | 1:205100172 | 0 | frameshift |  |  |  |  |
| c.762G>A;<br>p.W254* | 1:205100055 | 0 | nonsense |  |  |  |  |
| c.888A>T;<br>p.E296D | 1:205099929 | 0 | missense | Probably<br>damaging | 0.401 | 13.46 |  |
| c.919C>T;<br>p.R307* | 1:205099799 | 0 | nonsense |  |  |  |  |
| c.1490C>G;<br>p.S497C | 1:205094971 | 2.40e-6 | missense | Benign | 0.124 | 23.1 |  |
| c.1589-3T>C | 1:205088818 | 7.010e-7 | intronic |  |  |  | 0.06 |

**Supplemental Figure 1.**
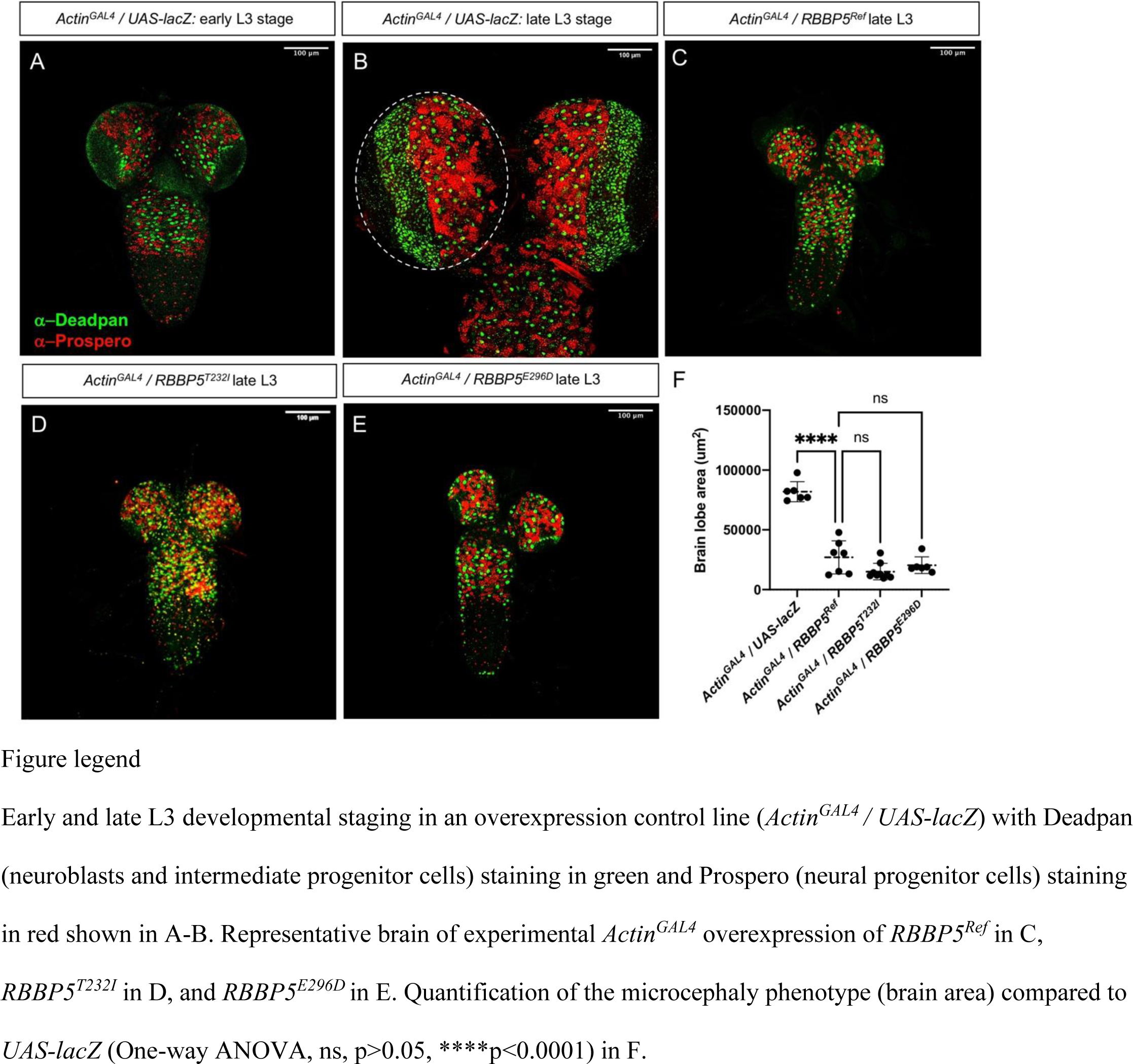
Human *RBBP5* transgenes interfere with *Drosophila* brain development.

**Supplemental Figure 2.**
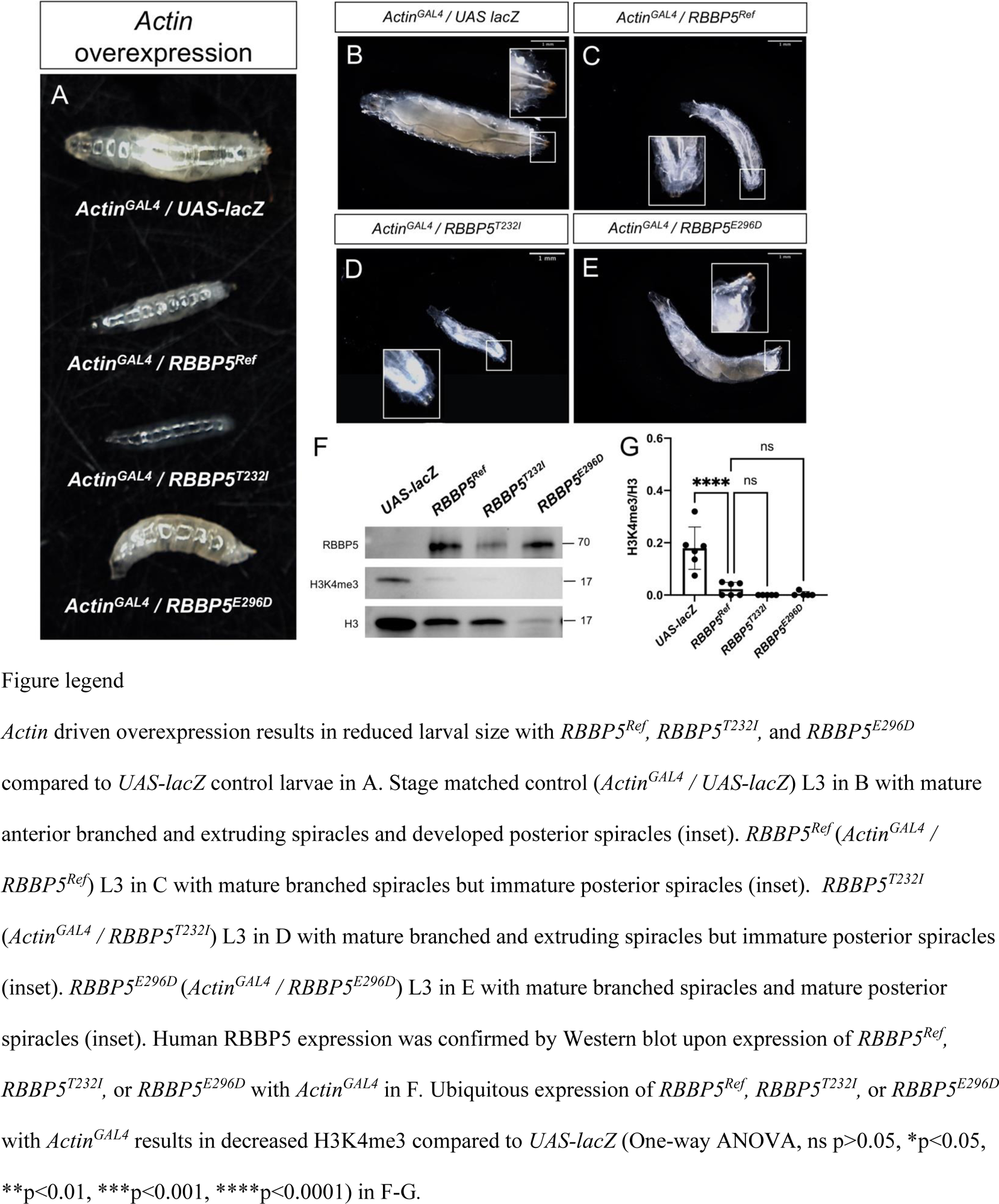
R*B*BP5 expression impairs L3 developmental progression and results in severely reduced H3K4 trimethylation.

**Supplemental Figure 3.**
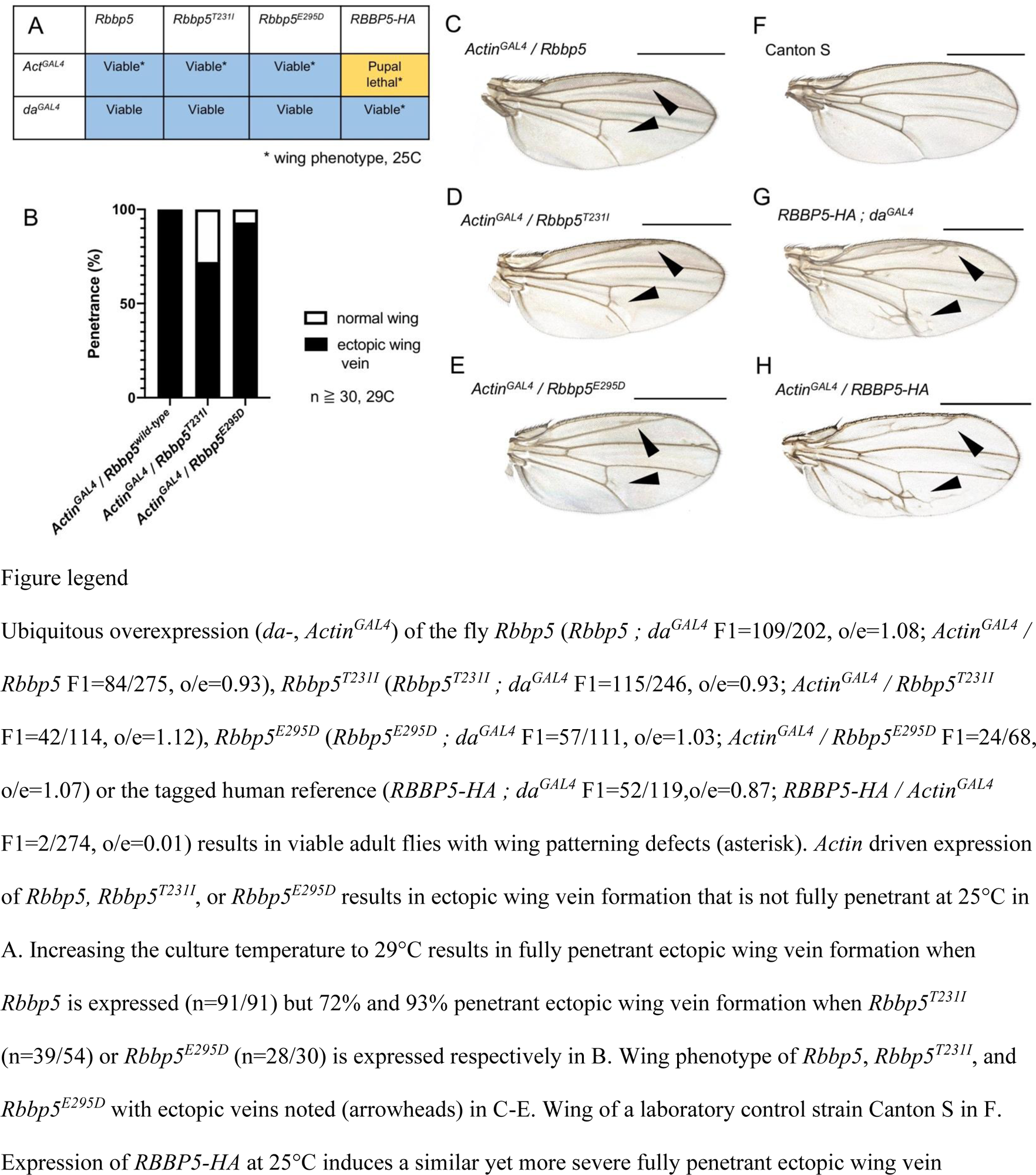
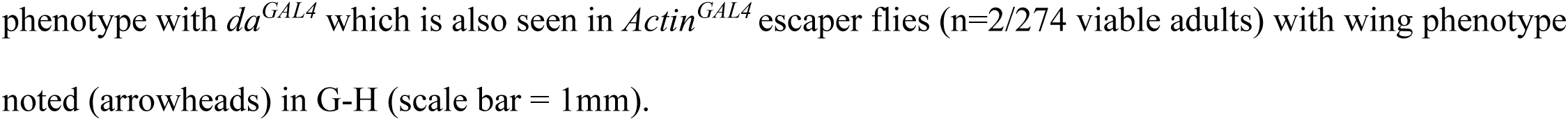
Overexpression of *Rbbp5/RBBP5-HA* induces ectopic wing vein formation.

